# Bioorthogonal Catalytic Microneedles Based on a Cytotoxic PEI Matrix for Synergistic Melanoma Therapy

**DOI:** 10.64898/2026.03.30.715245

**Authors:** Qian-He Xu, En-Kui Huang, Ya-Jun Chu, Xiaojun Yao, Pei-Nian Liu

**Affiliations:** Shanghai Key Laboratory of Functional Materials Chemistry, Key Laboratory for Advanced Materials, School of Chemistry and Molecular Engineering, East China University of Science & Technology, Shanghai 200237, China; State Key Laboratory of Natural Medicines, School of Pharmacy, China Pharmaceutical University, Nanjing 211198, China; Centre for Artificial Intelligence Driven Drug Discovery, Faculty of Applied Sciences, Macao Polytechnic University, Macao 999078, China; State Key Laboratory of Neurology and Oncology Drug Development, Nanjing, China

**Keywords:** Bioorthogonal catalysis, Microneedles, Polyethyleneimine, Synergistic therapy, Prodrug activation, Melanoma

## Abstract

Microneedle (MN) patches have emerged as a highly efficient platform for localized drug delivery, showing great promise in cancer therapy due to their ability to enable precise drug administration. However, conventional MN systems are limited by the low drug-loading capacity of their tips and primarily rely on biologically inert, non-therapeutic matrices for structural support, which restricts further gains in antitumor efficacy. Herein, we present a strategy turning toxicity into therapy by constructing palladium nanoparticle-loaded polyvinyl alcohol/polyethyleneimine (PVA/PEI@Pd) hydrogel microneedles (PPPd-MNs), which exploit the intrinsic cytotoxicity of PEI for synergistic melanoma therapy. The PPPd-MNs efficiently catalyze the deprotection of a doxorubicin prodrug (P-DOX), enabling *in situ* generation of active doxorubicin (DOX). Notably, the PEI matrix serves a dual function: acting as a robust ligand to stabilize Pd catalysts and functioning as a therapeutic agent that disrupts cancer cell membranes. Both *in vitro* and in *vivo* experiments demonstrate that the combination of Pd-mediated bioorthogonal activation of DOX and PEI-induced membrane damage achieves a remarkable synergistic therapeutic outcome in a murine melanoma model, resulting in a tumor inhibition rate of up to 98%. This work repurposes the inherent cytotoxicity of the carrier material as an active therapeutic component, offering a novel paradigm for the design of high-performance bioorthogonal catalytic systems.

## 1. INTRODUCTION

Microneedle (MN) patches represent an advanced transdermal delivery platform that enables efficient transport of drugs or biotherapeutics by painlessly penetrating the stratum corneum barrier via an array of micrometer-scale needles.^1–3^ Over the past decades, diverse MN formulations have been engineered to address specific biomedical needs, evolving from early solid^4^ and coated^5^ designs to increasingly sophisticated hollow,^6, 7^ dissolving,^8, 9^ and hydrogel-forming^10, 11^ systems. In cancer therapy, MNs offer a distinct advantage for localized therapy by enabling targeted delivery^12, 13^ or catalytic release^10, 11^ of chemotherapeutics or immunomodulators to superficial tumor sites, thereby minimizing systemic side effects associated with conventional chemotherapy. However, a common limitation in current tumor-treating MN systems is the low drug-loading capacity of the needle tips, which often leads to suboptimal therapeutic outcomes. The tips of microneedle systems typically consist of a polymeric matrix—constituting the bulk of the structure—and the loaded therapeutic agent. To date, these matrices have predominantly relied on biologically inert polymers, such as hyaluronic acid (HA),^12, 14^ polyvinyl alcohol (PVA),^10, 11^ polyvinylpyrrolidone (PVP),^15, 16^ or carboxymethyl cellulose (CMC).^17, 18^ These carriers typically serve merely as passive vehicles for drugs storage and release, lacking intrinsic therapeutic activity and thus contributing negligible antitumor effects on their own. Therefore, we envision using a structural matrix with inherent therapeutic toxicity in MN systems, which would amplify antitumor efficacy through synergistic mechanisms.

**Scheme 1.**
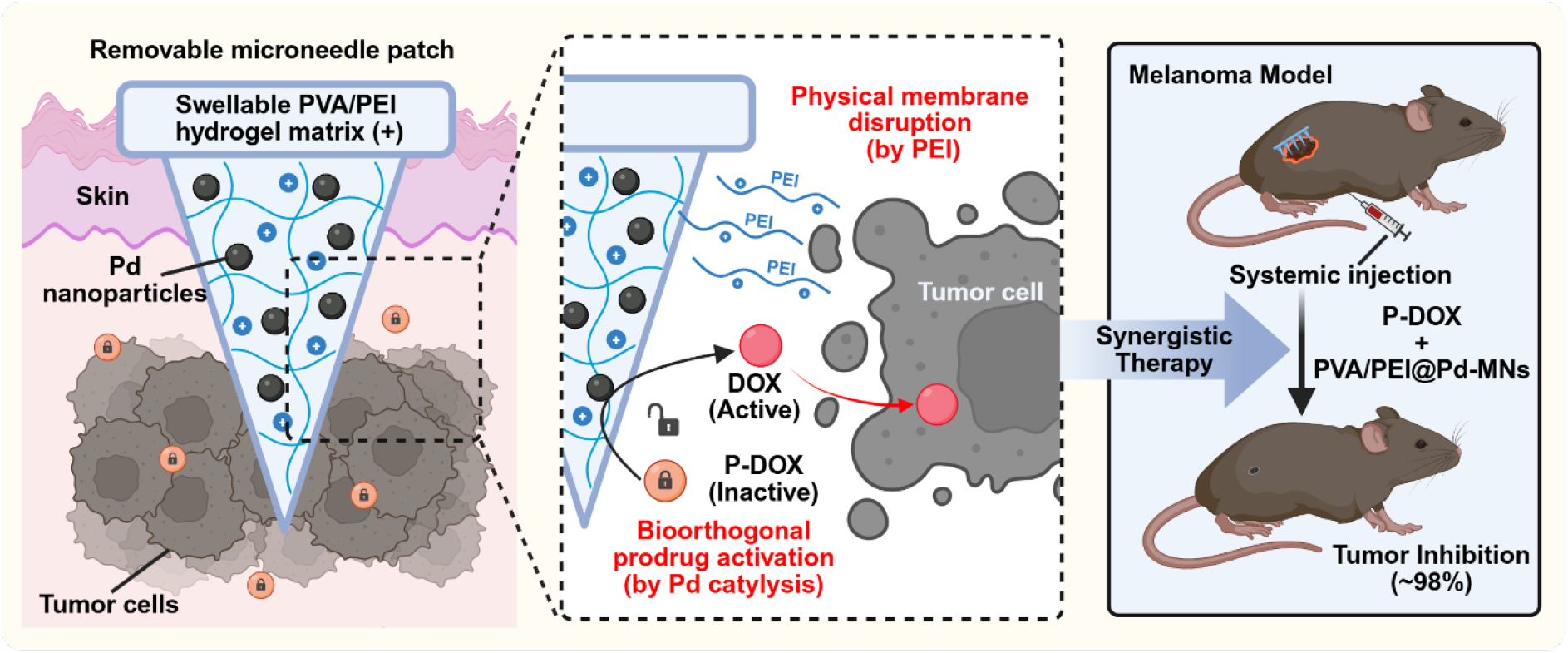
Schematic of bioorthogonal catalytic microneedles based on a cytotoxic PEI matrix for synergistic melanoma immunotherapy. PEI is incorporated into PVA matrix to form swellable hybrid hydrogel microneedle patches (PPPd-MNs). As a powerful ligand, PEI stabilized palladium nanoparticles and served as a therapeutic agent for physical destruction of tumor cell membranes. Meanwhile, the prodrug of doxorubicin is locally activated by the Pd catalyst in the swellable microneedle patch, directly killing the tumor.

Polyethyleneimine (PEI), a cationic polymer characterized by a high density of amine groups, has been extensively explored as a non-viral gene vector.^19, 20^ PEI exhibits pronounced cytotoxicity, primarily due to its inherently high cationic charge density, which facilitates strong electrostatic interactions with negatively charged cell membranes. This interaction destabilizes the membrane integrity, leading to physical disruption and subsequent necrosis or apoptosis of the cells.^21–23^ While this cytotoxicity of PEI is typically viewed as an adverse side effect to be mitigated in gene delivery, they present a unique opportunity for direct cancer cell eradication in localized therapies. Furthermore, the abundant amine groups in PEI possess strong coordination abilities with metal ions,^24–26^ making it an ideal candidate for anchoring and stabilizing transition metal nanoparticles.

Herein, we engineered a PEI-based hydrogel microneedle patch loaded with palladium nanoparticles (PPPd-MNs) to achieve synergistic therapy for melanoma. As illustrated in **Scheme 1**, PEI was incorporated into a PVA matrix to form a swellable hybrid hydrogel (PVA/PEI@Pd). This design endows the system with dual functionality: (1) serving as a robust ligand to stabilize Pd nanoparticles and prevent leakage; and (2) acting as a therapeutic agent to physically disrupt tumor cell membranes. Upon insertion into tumor tissue, the immobilized Pd nanoparticles catalyze the depropargylation of a doxorubicin prodrug (P-DOX), converting it into active doxorubicin (DOX) and thereby restoring its cytotoxicity. Simultaneously, the PEI matrix exerts cationic toxicity against cancer cells. Our results indicate that the combination of Pd-mediated bioorthogonal activation of chemotherapy and PEI-induced membrane damage achieves highly efficient tumor growth inhibition (~98%) in a mouse melanoma model. By transforming the toxic side effects of a carrier material into a synergistic therapeutic weapon, this work provides a novel therapeutic paradigm for the field of bioorthogonal catalysis.

## 2. MATERIALS SECTION

The detailed methods of the preparation of PVA/PEI@Pd and PPPd-MNs, characterizations, and the cellular and animal experiments are described in the Supporting Information.

## 3. RESULTS AND DISCUSSION

### 3.1. Preparation and Characterization of Catalytically Active PVA/PEI@Pd

Polyvinyl alcohol (PVA) was selected as the primary hydrogel matrix due to its excellent biocompatibility and ability to form suitable physical crosslinks with other polymers.^10, 11, 27, 28^ Polyethyleneimine (PEI), known for its chemical stability and strong interaction capabilities with polymers and metal ions,^25, 26, 29–31^ was physically crosslinked with PVA to serve as a swellable support for stabilizing palladium (Pd) nanocatalysts (**Figure 1a**). PdCl_4_^2-^ adsorbed within the PVA/PEI network was reduced to Pd nanoparticles using hydrazine hydrate. Fourier transform infrared (FTIR) spectroscopy confirmed the successful preparation of PVA/PEI@Pd (**Figure S1**). The spectrum exhibited a broad and intense peak at 3650–2991 cm^-1^, attributed to n(O-H) and n(N-H) stretching vibrations, and a n(C-O) stretching peak at 1085 cm^-1^. Notably, the retention of d(O-H) bending at 1659 cm^-1^, n(O-H) at 1563 cm^-1^, and d(-NH-) bending at 1454 cm^-1^ suggests a favorable chemical environment for Pd stabilization. Thermogravimetric analysis (TGA) (**Figure S2**) revealed that the onset decomposition temperature increased from 200 °C to 220 °C upon Pd loading. Furthermore, the residual weight above 500 °C significantly increased (approx. 15% for PVA/PEI@Pd vs. ~0% for PVA/PEI), indicating a systematic enhancement in thermal stability due to the incorporation of Pd nanoparticles.

**Figure 1.**
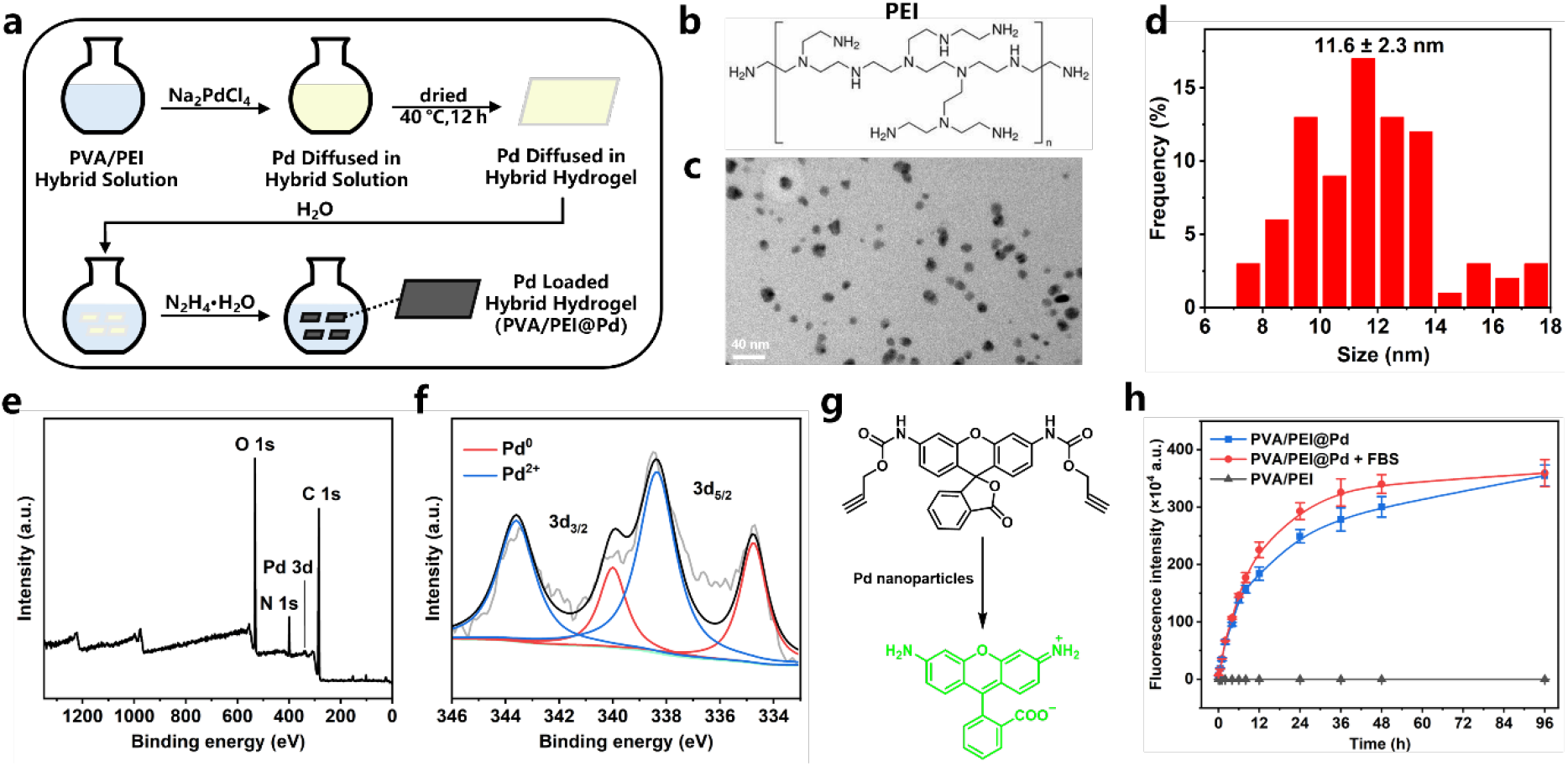
Preparation and characterization of the Pd loaded hybrid hydrogel (PVA/PEI@Pd). a) Schematic diagram for preparing PVA/PEI@Pd. b) Structure of PEI. c) TEM image of PVA/PEI@Pd. Scale bar, 20 nm. d) Diameter distribution histograms of Pd nanoparticles in (c). e) X-ray photoelectron spectrum (XPS) of PVA/PEI@Pd. f) Part of the XPS in (e). g) Schematic diagram depicting the decomposition of non-fluorescent P-Rho into fluorescent Rho 110. h) Changes in fluorescence intensity of the Rho 110 released by PVA/PEI@Pd in PBS buffer or PBS buffer containing 10% FBS. Data points represent means ± SD (n = 3) from three independent experiments.

Representative transmission electron microscopy (TEM) images showed that the reduced Pd nanoparticles within the PVA/PEI@Pd matrix had a diameter of 11.6 ± 2.3 nm (**Figure 1c-d**). Elemental mapping (**Figure S4**) confirmed the homogeneous distribution of Pd throughout the polymer matrix (**Figure S3**). X-ray photoelectron spectroscopy (XPS) was employed to investigate the chemical state of Pd (**Figure 1e**). Deconvolution of the Pd 3d peaks revealed a mixture of Pd(0) and Pd(II) states with a ratio of approximately 1:2 (**Figure 1f**). Inductively coupled plasma atomic emission spectroscopy (ICP-AES) analysis determined the metal content in the prepared PVA/PEI@Pd to be approximately 2.0 wt%. Given reports that both Pd(0) and Pd(II) can catalyze depropargylation reactions,^11, 32–35^ we synthesized a propargyloxycarbonyl-protected Rhodamine 110 (P-Rho) as a substrate to verify catalytic efficacy.

The prepared PVA/PEI@Pd was immersed in phosphate-buffered saline (PBS, pH 7.4) to reach swelling equilibrium, followed by the addition of the protected rhodamine for heterogeneous catalysis. The catalytic deprotection capability was revealed by monitoring the increase in green fluorescence (**Figure 1g**) and quantifying it against a standard curve (**Figure S5**). Time-dependent profiles (**Figure 1h**) demonstrated the effective uncaging of P-Rho by PVA/PEI@Pd. Notably, the catalytic performance in the presence of 10% fetal bovine serum (FBS) was superior to that in serum-free conditions. This enhancement may be attributed to the promoting effect of small molecular proteins (e.g., glutathione) in serum,^11, 32, 35, 36^ a characteristic advantageous for catalysis in biological environments. The above results all indicate the successful preparation of PVA/PEI@Pd with bioorthogonal catalytic activity.

### 3.2. Fabrication of Catalytically Active PPPd-MNs

Building on established protocols,^11^ PPPd-MNs were fabricated using polydimethylsiloxane (PDMS) molds via a dissolution-regeneration process of PVA/PEI@Pd. The resulting patch featured a 15 × 15 array of conical needles (**Figure 2a-c**) with a height of 1500 μm, a tip radius of 10 μm, a base diameter of 500 μm, and an inter-tip spacing of 800 μm. Scanning electron microscopy (SEM) of the longitudinal section (**Figure 2d** and **Figure S6**) confirmed the uniform distribution of Pd throughout the needle body. The mechanical strength of a single microneedle was measured to be 0.83 N (**Figure 2e**), showing no significant deviation from blank PVA/PEI microneedles. The incorporation of Pd nanoparticles did not compromise the mechanical integrity, likely due to the excellent dispersion of Pd and its interaction with the PVA/PEI network. This sufficient mechanical strength ensures effective skin penetration. The swelling behavior of PPPd-MNs was investigated by optical microscopy (**Figure 2f** and **Figure S7**). Upon immersion in PBS, the microneedles swelled rapidly, with increases in length and width of approximately 53% and 74%, respectively, within 30 minutes, demonstrating excellent swelling properties conducive to solute diffusion.

**Figure 2.**
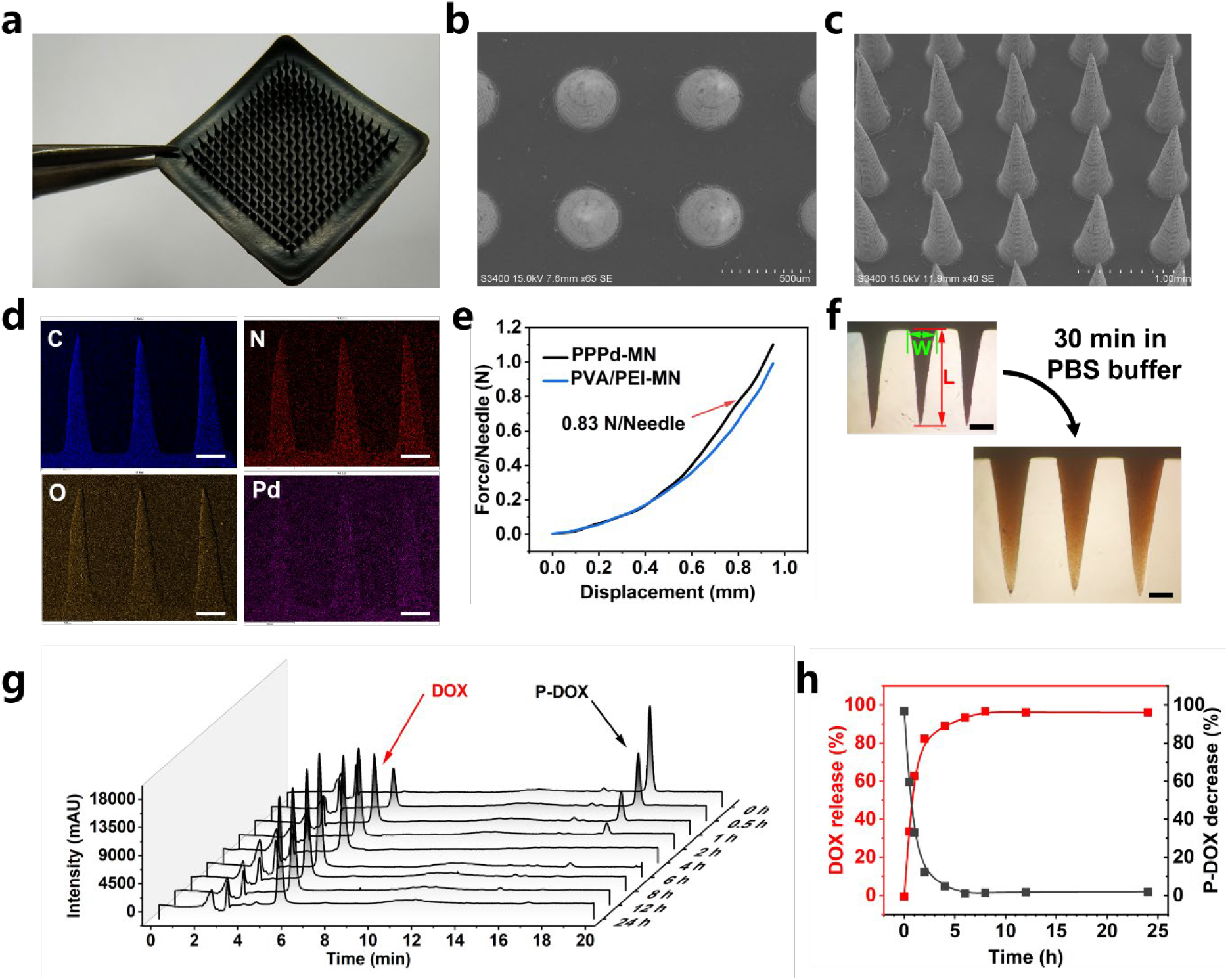
Preparation and characterization of PPPd-MNs. a) Photograph of a PPPd-MN. b, c) SEM top (b) and oblique (c) images of a PPPd-MN. Scale bar, 500 μm (b) and 1 mm (c). d) Elemental map analysis of a longitudinal section of a PPPd-MN. Scale bar, 300 μm. The energy dispersive X-ray spectrum was shown in **Figure S6.**e) The mechanical behaviors of PVA/PEI-MN and PPPd-MN. f) Photos of PPPd MNs soaked in PBS buffer for 0 min and 30 min. Scale bar, 300 μm. Evaluation of the swelling characteristics of PPPd-MNs in PBS buffer were shown in **Figure S7**. g) The catalytic release of PPPd-MNs on P-DOX over time measured by HPLC. h) The quantitative data of the P-DOX decrease and the DOX increase in (g).

### 3.3. PPPd-MN-Mediated Bioorthogonal Catalysis In Vitro

To evaluate the catalytic potential of the patch, we assessed the activation of a prodrug *in vitro*. A doxorubicin prodrug (P-DOX) was synthesized as a bioorthogonal candidate following reported procedures.^11^ Doxorubicin (DOX) acts by intercalating into DNA and inducing enzyme-mediated DNA damage;^37^ masking its amino group yields a prodrug with reduced activity that can be restored via bioorthogonal cleavage. Tests were conducted in a 3D-printed chamber designed to accommodate the patch submerged in 1 mL of PBS.^11^ High-performance liquid chromatography (HPLC) analysis indicated that PPPd-MNs catalyzed the deprotection of P-DOX in PBS, achieving a time-dependent release of active DOX with a conversion rate approaching 80% within 8 hours (**Figure 2g-h**). Catalyst leakage was assessed using ICP-AES, which detected only negligible Pd leakage (~1.1 wt%, 21.4 μg) into the solution after 24 hours. The stability of Pd within the matrix is likely ascribed to the strong coordination interactions between Pd and the PVA/PEI support.^29–31^

To further investigate bioorthogonal catalysis in a cellular environment, B16-F10 cells were cultured in medium containing P-Rho (10 μM) with PPPd-MNs immersed in the system. Confocal microscopy imaging revealed a time-dependent increase in intracellular fluorescence (**Figure S9a-b**), whereas no fluorescence recovery was observed in control groups treated with P-Rho alone (**Figure S9c-d**) or PPPd-MNs alone (**Figure S9e-f**). Flow cytometry results corroborated these findings, showing a progressive increase in fluorescence intensity in the co-treatment group (**Figure S9g-j**). These results confirm that PPPd-MNs can mediate the uncaging of P-Rho to release the fluorophore, which is subsequently taken up by cells. A similar trend was observed in 4T1 cells (**Figure S10**), confirming the universality of this catalytic activity across different cell lines.

### 3.4. PPPd-MN-Mediated Antitumor Activity In Vitro

PEI is a cationic organic polymer with one of the highest known charge densities and is widely studied as a non-viral gene vector.^19, 38^ Its cationic nature allows it to bind to anionic components of the cell membrane, leading to membrane destabilization and physical disruption, which triggers necrosis or apoptosis.^21, 22, 39^ Considering these properties, we evaluated the potential cytotoxicity of PPPd-MNs. Cytotoxicity assays revealed that PPPd-MNs exhibited significant dose-dependent toxicity towards both B16-F10 and 4T1 cells (**Figure 3a**). Cell viability dropped below 50% at dosages of 4 mg and 3 mg, respectively. Importantly, this toxicity was far lower towards human umbilical vein endothelial cells (HUVECs), indicating a degree of selectivity for cancer cells. Interestingly, control PVA/PEI microneedles without Pd (PP-MNs) exhibited even stronger toxicity, causing near-complete inhibition of B16-F10 and 4T1 cells at a dosage of just 2 mg (**Figure S11**). The observation that PP-MNs are more toxic than PPPd-MNs suggests that the coordination of Pd with PEI consumes a portion of the free amine groups, thereby reducing the effective cationic charge density on the polymer surface. This attenuation of toxicity is crucial, as it positions PPPd-MNs to act synergistically with the catalytically released drug rather than causing excessive non-specific damage.

**Figure 3.**
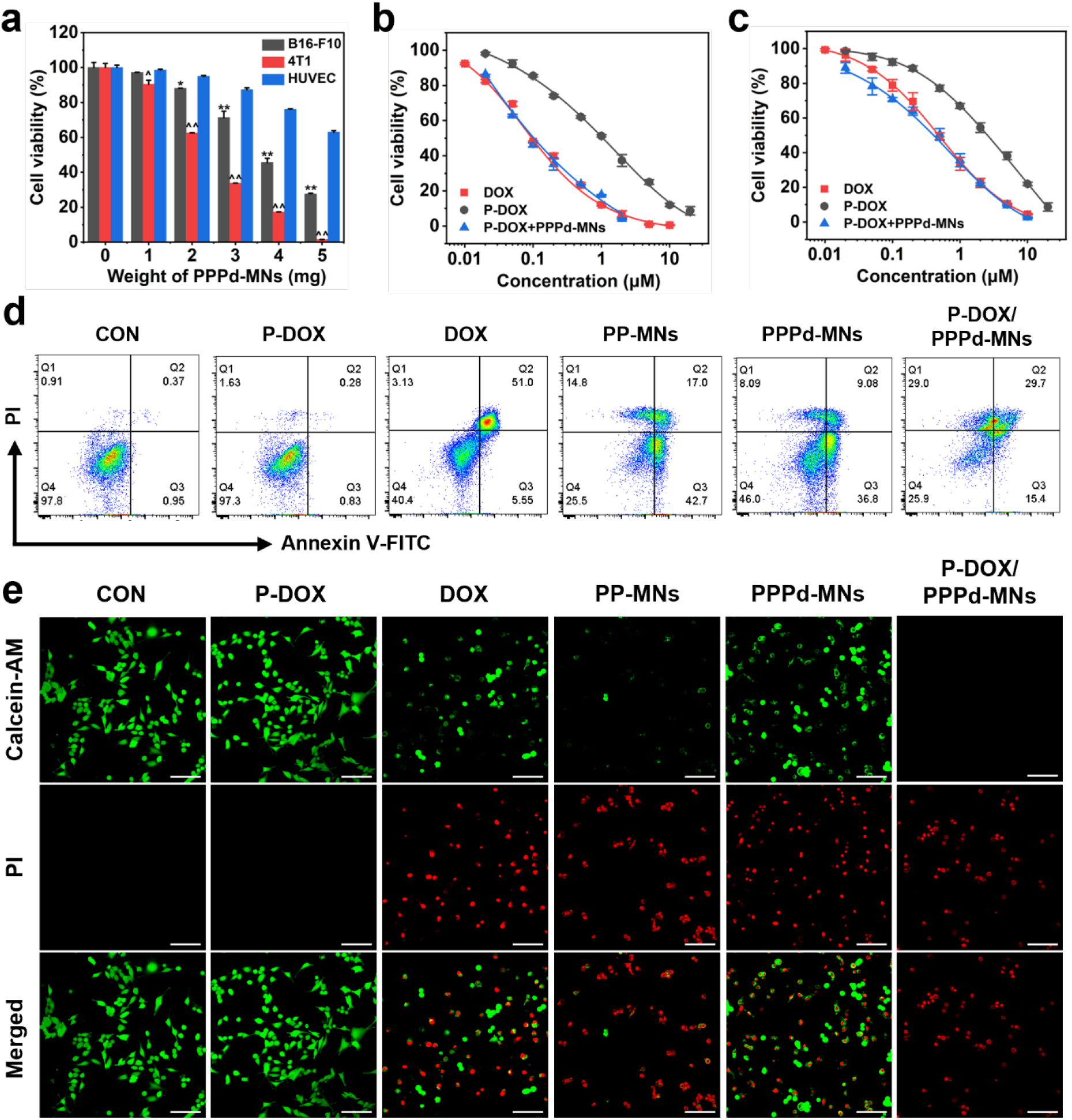
PPPd-MN-mediated antitumor activity *in vitro*. a) The cell viability of B16-F10, 4T1 and HUVEC cells treated with different dose of PPPd-MNs for 48 h. b, c) The cell viability of B16-F10 cells (b) and 4T1 cells (c) treated with different drug concentrations of DOX, P-DOX, and P-DOX/PPPd-MNs for 48 h. The data points in (a-c) represent the mean ± standard deviation (n = 3) from three independent experiments. d) Flow cytometry analysis of B16-F10 cell apoptosis after 24 hours of treatment with P-DOX, DOX, PP-MNs, PPPd-MNs or P-DOX/PPPd-MNs. e) Confocal laser microscopy image showing live and dead cell staining with calcein-AM and PI for B16-F10 cells, corresponding to the group shown in (d). Red fluorescence in the nucleus indicates cell death. Scale bar, 100 μm. *P < 0.05, **P < 0.01, ^P < 0.05 and ^^P < 0.01 vs. HUVEC groups in (a).

Melanoma is a highly malignant tumor known for its aggressive metastasis.^40^ To explore the prodrug activation performance mediated by PPPd-MNs, we selected a safe dosage of PPPd-MNs and evaluated the cytotoxicity of DOX, P-DOX, and the combination of P-DOX + PPPd-MNs against B16-F10 cells (**Figure 3b**). DOX exhibited potent cytotoxicity with increasing concentration (IC_50_ = 0.09 ± 0.02 μM), whereas P-DOX showed significantly lower toxicity (IC_50_ = 0.97 ± 0.12 μM). Notably, the P-DOX/PPPd-MNs group displayed an IC_50_ of 0.10 ± 0.02 μM, comparable to free DOX, indicating efficient prodrug activation. Similar results were obtained in 4T1 cells (**Figure 3c**), where the IC_50_ values for DOX, P-DOX, and P-DOX/PPPd-MNs were 0.45 ± 0.1 μM, 2.55 ± 0.3 μM, and 0.40 ± 0.08 μM, respectively.

We subsequently verified the synergistic effect between the intrinsic cytotoxicity of PPPd-MNs and bioorthogonal prodrug activation. Flow cytometry (**Figure 3d**) showed that B16-F10 cells treated with P-DOX/PPPd-MNs exhibited significant apoptotic and necrosis populations, exceeding that of the DOX-only or PPPd-MNs-only group. No obvious apoptotic cell population was detected in the P-DOX group. In addition, the PP-MNs group showed a certain trend of moderate apoptosis and necrosis, which was slightly attenuated in the PPPd-MNs group, consistent with the charge density hypothesis. Live/Dead staining (**Figure 3e**) further demonstrated massive cell death in the P-DOX/PPPd-MNs group, which was much greater than that of the DOX-only group. The PP-MNs group exhibiting a stronger trend of cell death compared to the PPPd-MNs group, while no significant apoptosis or cell death was observed in the control or prodrug groups. Similar to the cytotoxic activity experiment (**Figure S11**), the cell death observed in the MNs-only groups is likely driven by the inherent membrane-disrupting capability of PEI. These results confirm that PPPd-MNs not only effectively convert P-DOX into its active form but also exert intrinsic toxicity that synergizes with the chemotherapy.

Furthermore, ICP-AES analysis of the culture medium post-experiment detected negligible Pd leakage (~1.4 wt%), reinforcing the stability and biosafety of the system.

### 3.5. Synergistic In Vivo Melanoma Therapy Mediated by PPPd-MNs

Intracellular experiments suggested that the synergy between PPPd-MNs and bioorthogonal prodrug activation could benefit *in vivo* therapy. We first assessed the safety of systemic prodrug administration via intraperitoneal injection (i.p.), considering formulation and absorption.^11^ Mice were injected with varying doses of DOX or P-DOX every three days. P-DOX showed no adverse effects on body weight or major organ histology up to 100 mg/kg (**Figure S12-13**). In contrast, while 2.5 mg/kg DOX was safe, 5 mg/kg caused severe weight loss and cardiotoxicity (myocardial disarray and inflammation) (**Figure S13**). These data confirm that converting DOX to P-DOX significantly mitigates systemic toxicity, allowing for higher dosing.

Based on tolerance data, we utilized 50 mg/kg P-DOX for antitumor studies (20 times the safe dose of DOX) to maximize efficacy while avoiding side effects. The insertion capability of the MNs patch was verified (**Figure S14**), showing effective penetration and complete skin recovery within two hours post-removal. Patches were not reused due to being cut and minor deformation during application.

An orthotopic B16-F10 melanoma model was established in C57BL/6J mice to evaluate therapeutic efficacy. Mice were randomized into six groups: PBS, P-DOX, DOX, PP-MNs (catalyst-free control), PPPd-MNs, and P-DOX/PPPd-MNs. Treatments were administered every three days (two doses total), with patches removed 24 hours after each application. The P-DOX/PPPd-MNs group exhibited superior tumor suppression based on photos (**Figure 4a**) and tumor mass (**Figure 4d**). Tumor growth curves (**Figure 4b-c**) showed significant inhibition in the combination group with an inhibition rate of ~98%. The P-DOX group showed partial inhibition due to the high dosage permissible by its low toxicity. Notably, the PP-MNs and PPPd-MNs groups also exhibited tumor inhibition, validating the intrinsic therapeutic activity of the microneedle matrix itself. Consistent with *in vitro* data, the PPPd-MNs group showed slightly lower inhibition than PP-MNs, attributed to the reduced charge density of PEI upon Pd loading. H&E staining of tumor tissues (**Figure 4f**) confirmed extensive apoptosis in the P-DOX/PPPd-MNs group, surpassing that of DOX or PPPd-MNs alone. The therapeutic effect produced by the DOX or PPPd-MNs was not as good as that of the P-DOX/PPPd-MNs group, proving the synergistic therapeutic effect of the microneedle matrix and mediated bioorthogonal catalysis. The negligible body weight changes (**Figure 4e**) and histological analysis of major organs (**Figure S15**) attested to the biosafety of the therapy.

**Figure 4.**
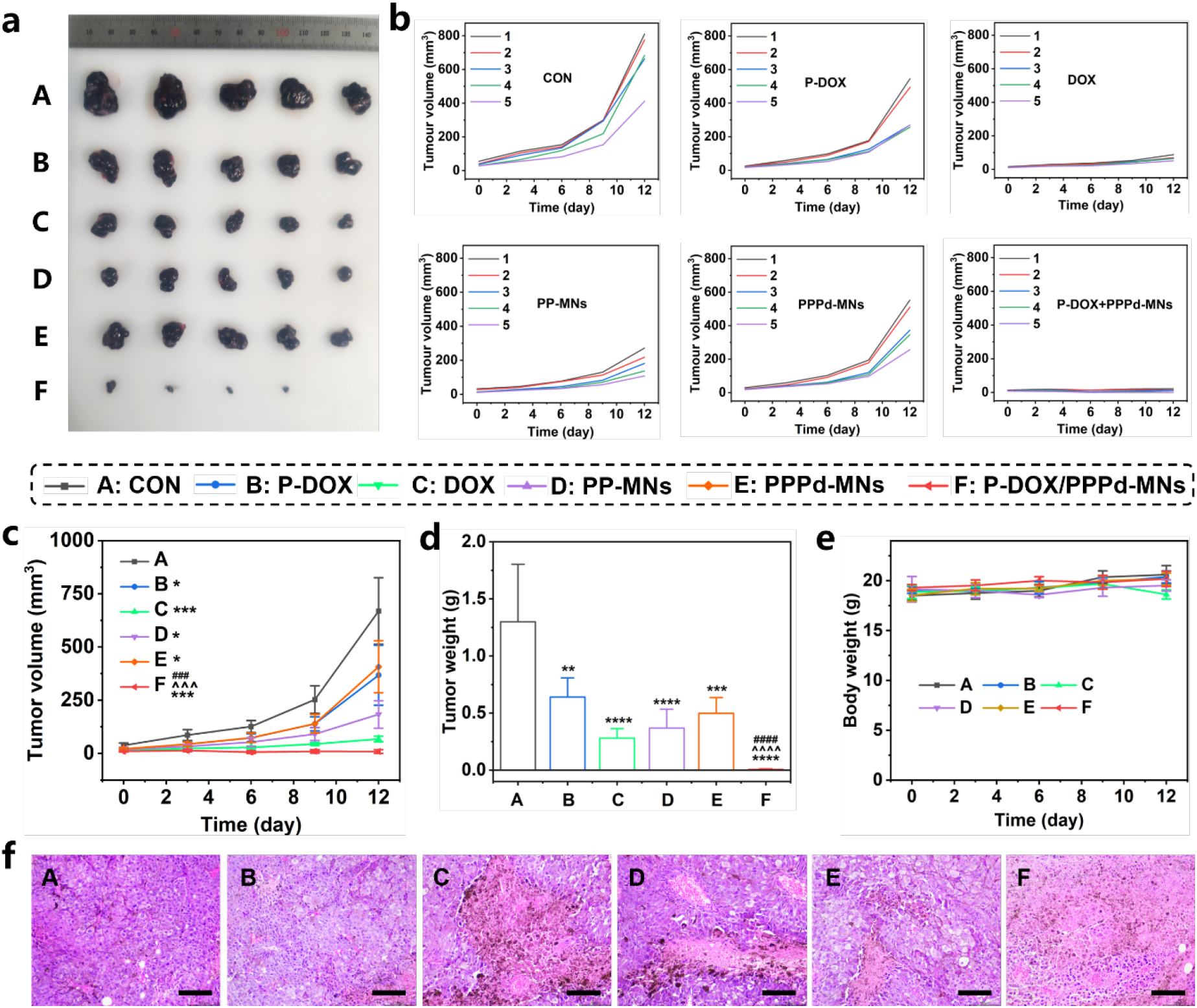
Synergistic *in vivo* melanoma therapy mediated by PPPd-MNs. a) Tumor photos of mice in different treatment groups after 12 days of treatment. b) Individual tumour growth kinetics of mice in each group. c) Average tumor growth curves of mice under different treatments. Data are shown as mean ± SD, n = 5. d) Average tumor weight of mice after 12 days of treatment. Data are shown as mean ± SD, n = 5. e) Body weight changes of mice during the observation period. Data are shown as mean ± SD, n = 5. f) Tumor tissue stained by H&E after 12 days of treatment. Scale bar = 100 μm. Data are shown as mean ± SD, n = 3. *P < 0.05, **P < 0.01, ***P < 0.001 and ****P < 0.0001 vs. control groups; ###P < 0.001 and ####P < 0.0001 vs. group C; ^^^P < 0.001 and ^^^^P < 0.0001 vs. group E.

Overall, the design of PPPd-MNs transcends the traditional limitation of bioorthogonal catalyst supports acting merely as inert scaffolds. Rather than solely pursuing biocompatibility, we strategically ‘tamed’ the high cationic toxicity of PEI through Pd coordination, modulating it to a balance point where it facilitates therapy via membrane disruption without causing uncontrollable toxicity. This feature combined with the *in situ* activated high-concentration chemotherapy, establishes a dual-strike mode of ‘physical membrane disruption + chemical DNA damage,’ thereby achieving superior antitumor efficacy compared to monotherapies. This synergistic strategy provides new insights for the development of multifunctional and integrated bioorthogonal catalytic devices.

## 4. CONCLUSION

In summary, we successfully developed the bioorthogonal catalytic microneedles (PPPd-MNs) based on a cytotoxic PEI matrix for the highly efficient synergistic treatment of melanoma. By incorporating PEI into the PVA microneedle matrix, we achieved not only the robust immobilization of Pd nanoparticles with minimal leakage but also creatively exploited the cationic nature of PEI to disrupt tumor cell membranes. This physical damage mechanism, combined with the Pd-catalyzed *in situ* activation of the P-DOX prodrug and generation of active DOX, led to a potent synergistic antitumor effect. Both *in vitro* and *in vivo* experiments confirmed that PPPd-MNs could effectively activate prodrugs to generate DOX and induce cancer cell death, achieving a tumor inhibition rate of up to 98% while maintaining biosafety. This work demonstrates the immense potential of transforming intrinsic material toxicity into synergistic therapeutic means, which provides a powerful tool for tumor-treating MN systems.

## ASSOCIATED CONTENT

### Supporting Information

The Supporting Information is available free of charge at DOI. FT-IR, TGA, TEM of PVA/PEI and PVA/PEI@Pd; Standard curve of Rho 110, DOX and P-DOX; SEM results of PPPd-MN; Swelling characteristics of PPPd-MNs; CLSM and cytotoxicity of P-Rho/PPPd-MNs on B16-F10 cells or 4T1 cells; The cell viability of B16-F10 cells and 4T1 cell treated with PPPd-MNs or PP-MNs; Assessment of systemic toxicity for DOX and P-DOX and other data (PDF)

## AUTHOR INFORMATION

### Author Contributions

This manuscript has been reviewed by all authors. All authors have given approval to the final version of the manuscript.

### Notes

The authors declare no competing financial interest.

## ACKNOWLEDGMENT

This work was supported by the National Natural Science Foundation of China (No. 92580139) and Macao Science and Technology Development Fund (0030/2024/RIA1). We thank the Research Center of Analysis and Testing of East China University of Science and Technology for help in the characterization.

