## Supplemental Information for "Bioorthogonal Catalytic Microneedles Based on a Cytotoxic PEI Matrix for Synergistic Melanoma Therapy"

### Experimental Section

#### Materials

All chemical reagents and solvents were purchased from Adamas-beta (Shanghai, China) and used as received unless otherwise noted. Ultrapure water (18.2 MΩ·cm$^{-1}$, Millipore) was used in all experiments. Silica gel for column chromatography (200-300 mesh) and silica gel plates for thin-layer chromatography (GF254) were purchased from Shanghai Zhonghe Chemical Technology Co., Ltd. (Shanghai, China). Polyvinyl alcohol (PVA, MW 89,000–98,000; degree of hydrolysis 99+%; Cat. No. 341584) was purchased from Sigma-Aldrich (St. Louis, USA). Dulbecco's Modified Eagle Medium (DMEM), Roswell Park Memorial Institute 1640 medium (RPMI 1640), and fetal bovine serum (FBS) were purchased from ThermoFisher Tech Co., Ltd. Cell Counting Kit-8 (CCK-8) was purchased from APExBIO (USA). The Annexin V-FITC/PI Apoptosis Detection Kit was purchased from Vazyme (Nanjing, China). Branched polyethyleneimine (PEI, MW 25,000; Cat. No. C0539) and the Calcein/PI Live/Dead Viability/Cytotoxicity Assay Kit were purchased from Beyotime (Nantong, China).

#### Characterization

$^{1}$H NMR spectra were recorded on a Bruker Advance 400 NMR spectrometer. Scanning electron microscopy (SEM) images were obtained using JSM-IT800HL and Hitachi S-3400N instruments. Transmission electron microscopy (TEM) images were acquired on a JEM-2100 electron microscope (accelerating voltage 200 kV). Fourier transform infrared (FTIR) spectra were recorded by a Nicolet 6700 spectrometer. Thermogravimetric analysis (TGA) curves were recorded by a TGA 8000 thermogravimetric analyzer. Fluorescence measurements were performed using a FluoroMax-4 spectrofluorometer. X-ray photoelectron spectroscopy (XPS) was determined using a Thermo Scientific K-Alpha instrument. Palladium (Pd) content analysis was completed using an Agilent 725 inductively coupled plasma atomic emission spectrometer (ICP-AES). The mechanical behavior of microneedles was analyzed by a universal testing machine (ZQ-990B, Zhiqu, Dongguan, China). Cell viability analysis was detected using a Synergy H4 microplate reader. Flow cytometry data were acquired via a Beckman CytoFLEX S flow cytometer. Confocal microscope images were taken using a confocal laser scanning microscope (STELLARIS 8, Leica, Germany).

**Preparation and Characterization of PVA/PEI@Pd**

PVA (4.8 g) was dissolved in 24 mL of deionized water at 95 °C with stirring. After cooling to 60 °C, PEI (1.2 g, MW=25000) was added and stirred for 30 min. To this mixture, 60 mL of freshly prepared $Na_2PdCl_4$ (20 mM) solution was added, and the mixture was stirred at room temperature in the dark for 2 h. The mixed solution was poured onto a glass plate to form a film and dried at room temperature. The prepared polymer film was immersed in 200 mL of 10% hydrazine hydrate solution for reduction for 2 h, washed with water three times (1 h each), and dried to obtain PVA/PEI@Pd. The prepared PVA/PEI@Pd and a PVA/PEI control film prepared in the same manner were used for FTIR and TGA testing. The PVA/PEI@Pd was re-dissolved in pure water and dropped onto a copper grid for TEM imaging and mapping analysis. The PVA/PEI@Pd was ground into small pieces of ~2 mm for XPS analysis. Inductively coupled plasma atomic emission spectroscopy (ICP-AES) analysis indicated that the palladium loading in PVA/PEI@Pd was ~2.0 wt%.

### Synthesis of P-Rho

The propargyl-protected probe P-Rho was synthesized according to previously reported work.[1]

### Depropargylation Mediated by PVA/PEI@Pd

PVA/PEI@Pd (1 mg) was added to PBS buffer (pH 7.4), followed by the addition of P-Rho (100 μL, 1 mM in DMSO) to reach a total system volume of 1 mL. For the serum group, the mixture contained 10% fetal bovine serum (FBS). The reaction mixture was stirred at 37 °C, and 20 μL samples were taken at different time points. Each sample was mixed with 780 μL of water, and the fluorescence intensity of Rho 110 was measured using fluorescence spectrophotometry (Excitation wavelength Ex = 488 nm, Emission wavelength Em = 530 nm) to monitor the reaction progress. Quantitative analysis was performed using the standard curve of Rho 110 (shown in **Figure S5**).

### Preparation of PPPd-MNs

This study prepared microneedles referring to a previously reported method.[1] PDMS silicone molds were purchased from Taizhou Weixin Pharmaceutical Technology Co., Ltd. The negative molds were formed by pouring PDMS into 3D-printed positive molds. The mold featured a 15 × 15 needle array with a pitch of 800 μm. Each needle cavity was conical, with a base diameter of 500 μm, a height of 1,500 μm, and a tip radius of 10 μm. PVA/PEI@Pd was mixed with water and heated to 95 °C to prepare a 15 wt% solution. A uniform mixture (0.8 mL) was carefully injected into the pre-wetted PDMS mold channels, avoiding air bubbles. The mold was dried at 25 °C for 48 hours without centrifugation or freezing. The microneedle array patch was carefully separated and stored in a desiccator for later use. PVA/PEI microneedles were prepared using the same 15% PVA/PEI solution. To characterize the distribution of Pd nanoparticles in the polymer matrix, the microneedles were embedded in EPON 812 resin, sectioned using a Leica EM UC7 ultramicrotome, and then subjected to SEM Mapping imaging.

**Synthesis of P-DOX**

The propargyl-protected doxorubicin prodrug P-DOX was synthesized according to previously reported work.[1]

**Prodrugs Activation Experiment**

Based on previously reported parameters, a batch of chambers with an internal volume of approximately 1 mL suitable for microneedle array patches was produced by 3D printing using biocompatible PolyJet material MED 610 (Stratasys).[1] The catalytic study of the microneedles was performed in a chamber with a total volume of 1 mL containing PBS buffer (pH 7.4) and P-DOX (100 μL, 1 mM in DMSO). The microneedle array patch was placed into the solution, covered with a lid, and placed in a 37 °C incubator. At designated time points, 10 μL of the reaction solution was removed and mixed with 90 μL of methanol for HPLC analysis. HPLC analysis was performed using a Shimadzu LC-20A system equipped with a WondaSil C18 Superb reversed-phase column (particle size 5 μm, 4.6 × 150 mm), with a flow rate of 1.0 mL/min and a column temperature of 30 °C. The mobile phase gradient for the P-DOX reaction solution was as follows: 0.00-5.30 min, water/acetonitrile (70:30); 5.31-17.00 min, water/acetonitrile (58:42); 17.00-20.00 min, water/acetonitrile (70:30). The aqueous phase contained 0.1% trifluoroacetic acid. All chromatograms were detected at a wavelength of 254 nm, and prodrug conversion and drug release rates were calculated using standard curves (shown in **Figure S8**).

**Catalyst Leakage studies**

To test the stability of PPPd-MNs against palladium leakage, reaction solutions or cell culture media involving PPPd-MNs were collected, concentrated, and digested with concentrated acid, followed by analysis via inductively coupled plasma atomic emission spectroscopy (ICP-AES).

**Cell culture**

B16-F10 and 4T1 cells were purchased from Shanghai Institute of Cells, Chinese Academy of Sciences. B16-F10 cells were cultured in RPMI 1640 medium, and 4T1 cells were cultured in DMEM. Both media were supplemented with 10% (v/v) fetal bovine serum (GIBCO) and 1% penicillin-streptomycin (GIBCO, Invitrogen). All cells were cultured in a 37 °C, 5% $CO_2$ incubator (Thermo Scientific).

**PPPd-MNs-Mediated Bioorthogonal Catalytic Reaction in Cell Culture**

The degradation of P-Rho by PPPd-MNs in cell culture was investigated by confocal microscopy and flow cytometry. For confocal microscopy analysis, B16-F10 or 4T1 cells were seeded at a density of $5 \times 10^4$ cells/well in 3D-printed culture chambers with glass coverslips at the bottom. After 24 hours, the medium was replaced with fresh medium, and the following treatments were applied at different time points: 10 μM P-Rho combined with PPPd-MNs, 10 μM P-Rho alone, or PPPd-MNs alone. The PPPd-MNs were placed in the culture chamber so that the microneedles were immersed in the medium. Nuclei were stained with Hoechst 33342, and imaging was performed through the glass coverslip using a confocal microscope. For flow cytometry analysis, cells in each culture chamber were resuspended, and 10,000 events per sample were analyzed using an A525/40 bandpass filter (FITC).

**In Vitro Cytotoxicity Assays**

B16-F10, 4T1, or HUVEC cells were seeded in 24-well plates at a density of $5 \times 10^4$ cells/well. After culturing for 24 hours, cut microneedles of different masses were added (n=3 replicates per group), with a medium volume of approximately 500 μL per well. The control group received PP-MNs prepared in the same manner. After continuing culture for 48 hours, cell viability was detected using the standard Cell Counting Kit-8 (CCK-8) method, measuring absorbance at 450 nm using a microplate reader. For the P-DOX/PPPd-MNs group, a fixed mass (1 mg) of PPPd-MNs was immersed in the medium after adding the prodrug. To analyze potential palladium leakage, the medium was transferred to a centrifuge tube, digested with concentrated nitric acid, diluted, and detected using ICP-AES.

**In Vitro Apoptosis Assay**

B16-F10 cells were seeded in 6-well plates at a density of $2.5 \times 10^5$ cells/well. After culturing for 24 hours, cells were treated with 2 μM P-DOX, DOX, PP-MNs, PPPd-MNs or P-DOX/PPPd-MNs for 24 hours. The control group received an equivalent amount of DMSO (maximum concentration). After treatment, cells were double-stained with Annexin V-FITC and propidium iodide (PI) according to the manufacturer's instructions, and apoptosis was detected by flow cytometry.

**In Vitro Live/Dead Staining Assay**

The Calcein-AM/PI double staining method was used to further evaluate the therapeutic effect of PPPd-MNs combined with P-DOX. B16-F10 cells were seeded in confocal dishes at a density of $2 \times 10^5$ cells/dish. After culturing for 24 hours, cells were treated with 5 μM P-DOX, DOX, PP-

MNs, PPPd-MNs or P-DOX/PPPd-MNs for 24 hours. The control group received an equivalent amount of DMSO (maximum concentration). After treatment, cells were washed with PBS and incubated with Calcein-AM and PI solution for 30 minutes to label live and dead cells. After washing again with PBS, imaging was performed using confocal laser scanning microscopy (CLSM). Live cells were labeled with green fluorescence by Calcein-AM, and dead cells were labeled with red fluorescence by PI, allowing for the assessment of cell survival and the therapeutic efficacy of the treatment regimens.

**Animal Experiments**

Healthy female C57BL/6 mice (6-8 weeks) were purchased from GemPharmatech Co., Ltd. (Shanghai, China). All animal experiments involved randomly grouping the mice. A solid tumor model was established by subcutaneously injecting B16-F10 melanoma cells into the ventral side of C57BL/6J mice. All drugs were administered intraperitoneally every three days for a total of two injections. For the PPPd-MNs and PP-MNs treatment groups, the PPPd-MNs or PP-MNs were resized to cover the tumor area and fixed with a waterproof and breathable polyurethane film to ensure full swelling of the microneedles. All animal experiments followed the Guidelines for the Care and Use of Laboratory Animals and were approved by the Institutional Animal Care and Use Committee (IACUC, Approval No.: SHRM-IACUC-089).

**Drug Side Effect in Vivo**

DOX and P-DOX solutions were prepared by dissolving in PBS buffer supplemented with 2% Tween-80. To determine a suitable dose for anti-tumor studies, the toxicity of P-DOX was evaluated by measuring body weight changes in mice after intraperitoneal injection every three days. The physiological signs of DOX-treated mice were monitored for comparison. At the end of the experiment, drug side effects were assessed via body weight changes and H&E staining of various organs.

**In Vivo Antitumor Activity Mediated by PPPd-MNs Synergistic Bioorthogonal Catalysis**

B16-F10 melanoma cells ($1 \times 10^6$) were subcutaneously injected into the right flank of C57BL/6J mice. Seven days later, the mice were randomly divided into 6 groups: (a) PBS control group; (b) P-DOX group; (c) DOX group; (d) PP-MNs group; (e) PPPd-MNs group; (f) P-DOX/PPPd-MNs group. The PBS, DOX (2.5 mg $kg^{-1}$), and P-DOX (50 mg $kg^{-1}$) groups received intraperitoneal injections every three days, for a total of two times. The PPPd-MNs and PP-MNs groups had the patches resized to cover the tumor area, fixed with waterproof and breathable PU film, and removed after 24 hours. Tumor size was measured using a digital caliper, and tumor volume ($mm^3$) was calculated using the formula: long diameter × short diameter$^2$/2. To evaluate potential toxicity, mouse body weight changes were recorded. At the end of the treatment, tumors and major organs (heart, liver, spleen, lung, kidney) were subjected to Hematoxylin and Eosin (H&E) staining for histological analysis.

**Statistical Analysis**

Data in this study are expressed as mean ± standard deviation (SD). Statistical analysis was performed using Origin 10.0 software. All data were derived from three independent experiments. Significant differences were evaluated by two-tailed unpaired t-test, one-way analysis of variance (ANOVA), or repeated measures ANOVA. Significance thresholds were set at * $p < 0.05$, ** $p < 0.01$, *** $p < 0.001$, and **** $p < 0.0001$.

### Supplementary Figures

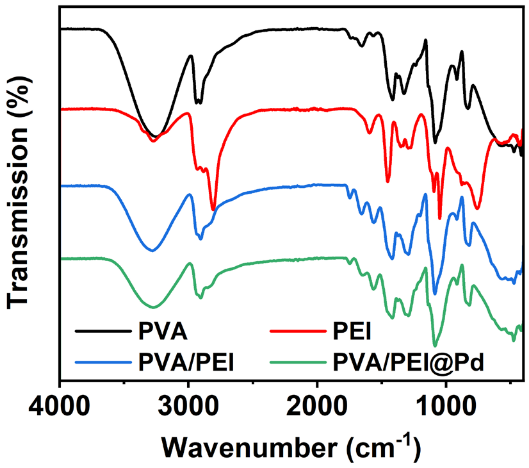


**Figure S1.** Frontier transform infrared spectra of PVA, PEI, PVA/PEI and PVA/PEI loaded with Pd nanoparticles (PVA/PEI@Pd).

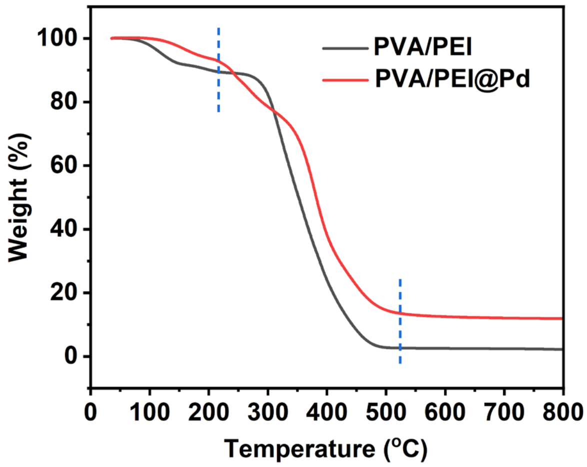


**Figure S2.** Thermogravimetric analysis (TGA) of PVA/PEI before and after Pd nanoparticles loading.

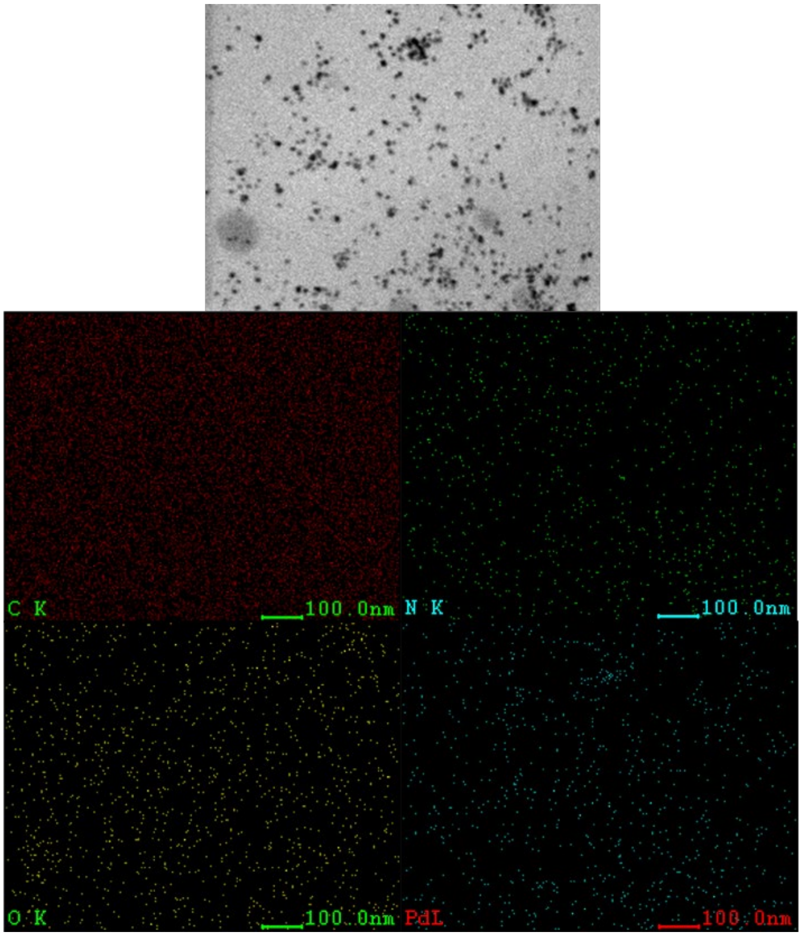


**Figure S3.** TEM image and elemental spectrum analysis of PVA/PEI@Pd. Scale bar: 100 nm.

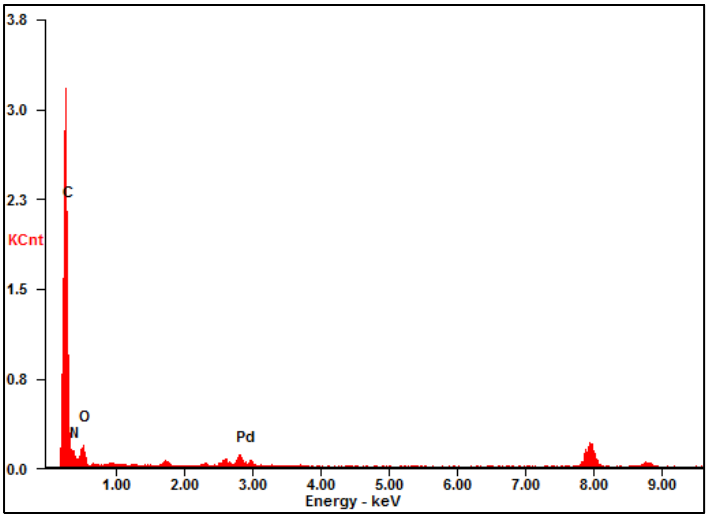


**Figure S4.** The energy dispersive X-ray spectrum of PVA/PEI@Pd corresponds to the elemental mapping analysis shown in **Figure S3.**

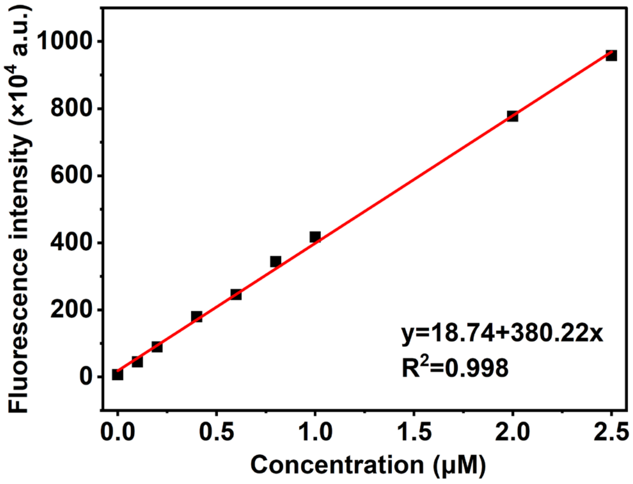


**Figure S5.** Linear fitting standard curve showing fluorescence intensity at 530 nm (Ex = 488 nm, Em = 530 nm) as a function of Rho 110 concentration. Data points represent means ± SD (n = 3) from three independent experiments.

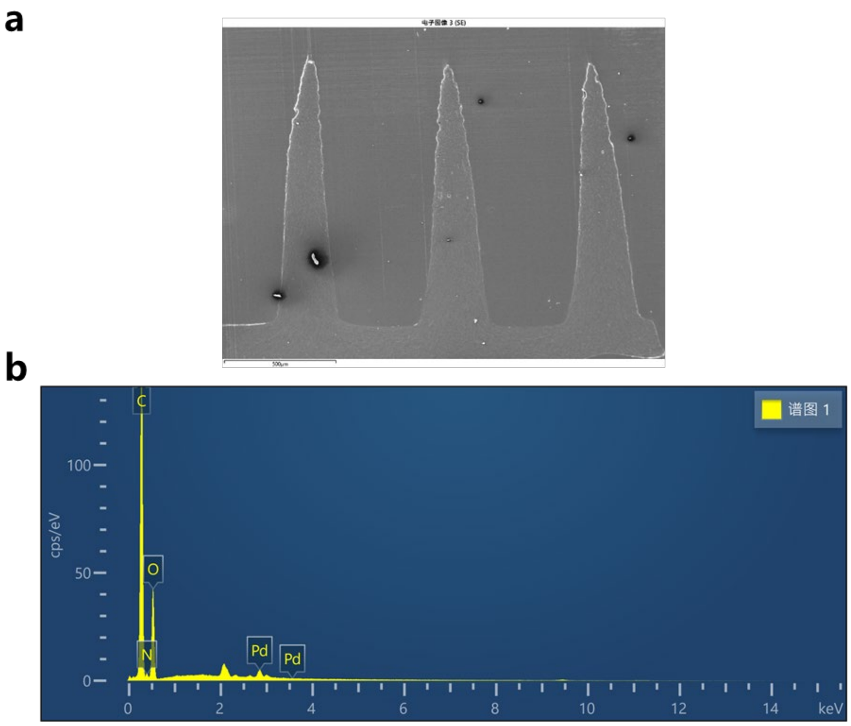


**Figure S6.** (a) SEM image of a longitudinal section of a PPPd-MN (scale bar, 500 μm). (b) The energy dispersive X-ray spectrum corresponds to the elemental mapping analysis of the microneedles cross-section shown in (a) and **Figure 2d**.

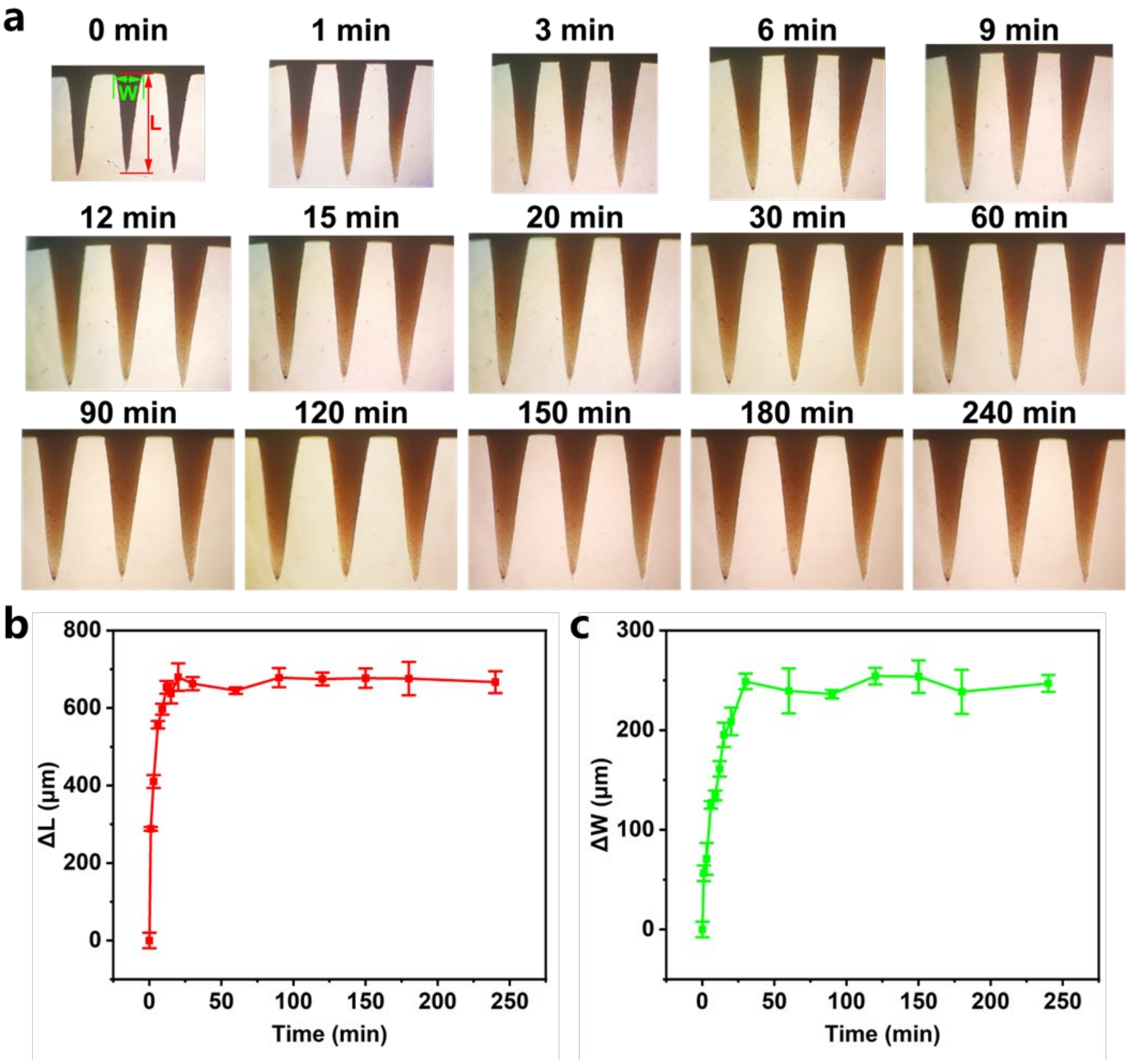


**Figure S7.** Evaluation of the swelling characteristics of PPPd-MNs in PBS buffer. (a) Representative optical micrographs displaying the morphological changes of microneedles at various time intervals following immersion in PBS buffer. Quantitative analysis of the variations in needle length (b, ΔL) and base width (c, ΔW) over time. The specific dimensions measured are indicated by the schematic inset in panel (a). Data are presented as mean ± SD (n = 3 independent experiments).

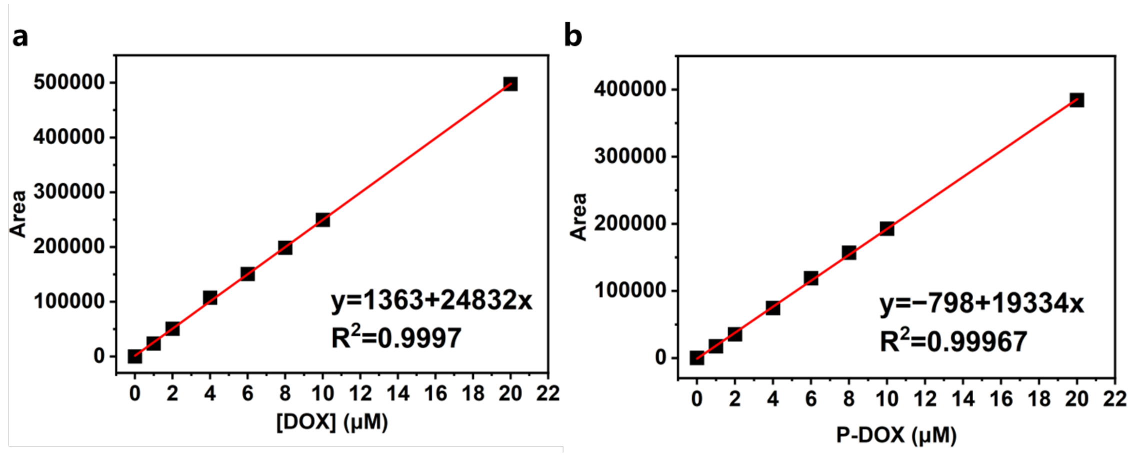


**Figure S8.** Linear fitting standard curve showing strong UV absorption of (a) DOX and (b) P-DOX detected by HPLC, which were used to calculate the conversion rate in **Figure 2h**.

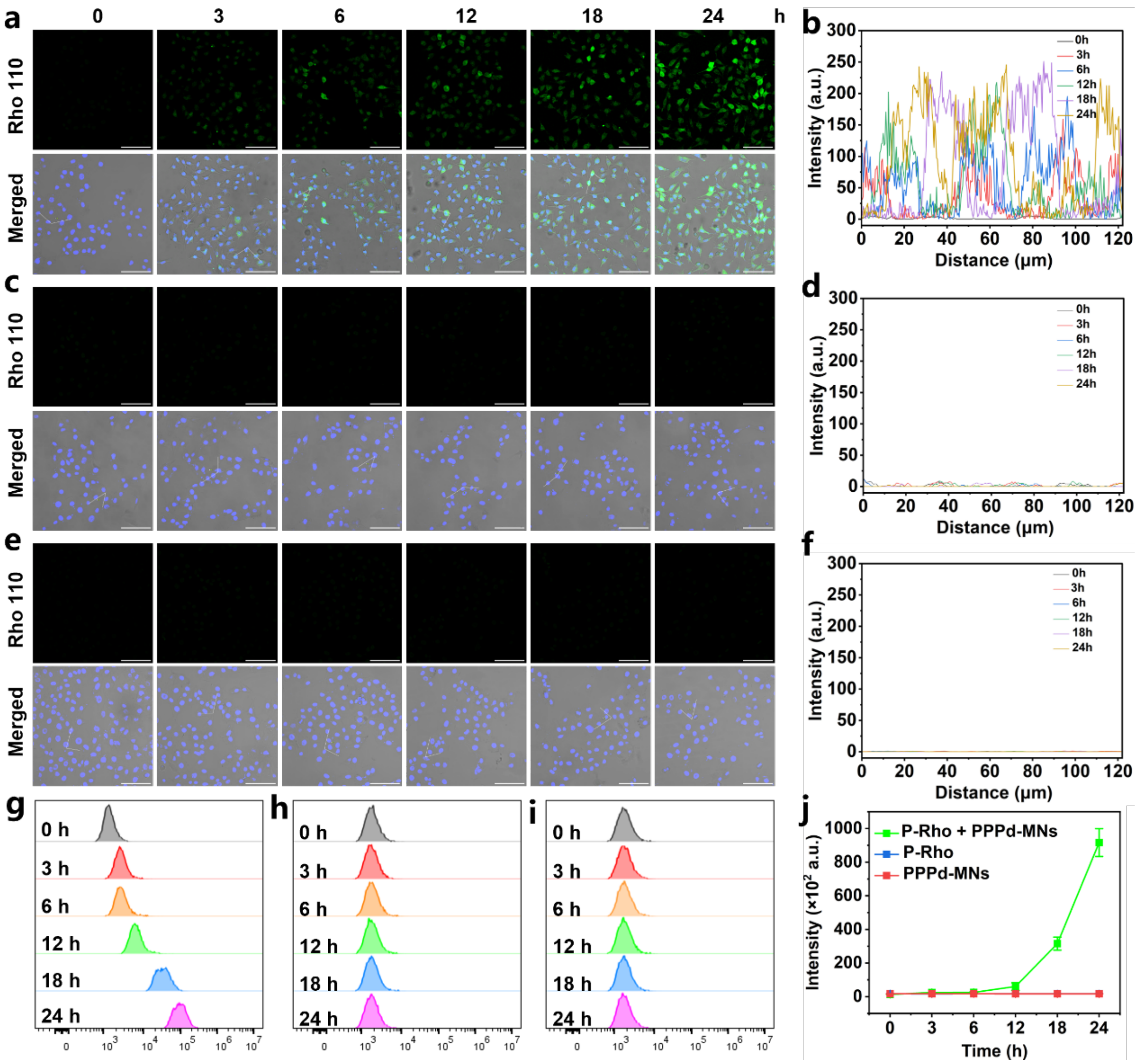


**Figure S9.** Intracellular bioorthogonal activation of P-Rho mediated by PPPd-MNs in B16-F10 cells. (a–f) Representative confocal laser scanning microscopy (CLSM) images visualizing the time-dependent fluorescence recovery in B16-F10 cells treated with the combination of P-Rho and PPPd-MNs (a, b), P-Rho alone (c, d), or PPPd-MNs alone (e, f). The corresponding line scan profiles (right panels) depict the fluorescence intensity distribution along the white dashed lines. Scale bar: 50 μm. (g–i) Flow cytometric histograms of B16-F10 cells subjected to different treatments over time: P-Rho + PPPd-MNs (g), P-Rho (h), and PPPd-MNs (i). (j) Quantified mean fluorescence intensity (MFI) derived from the flow cytometry data in panels (g–i). Data represent mean ± SD (n = 3).

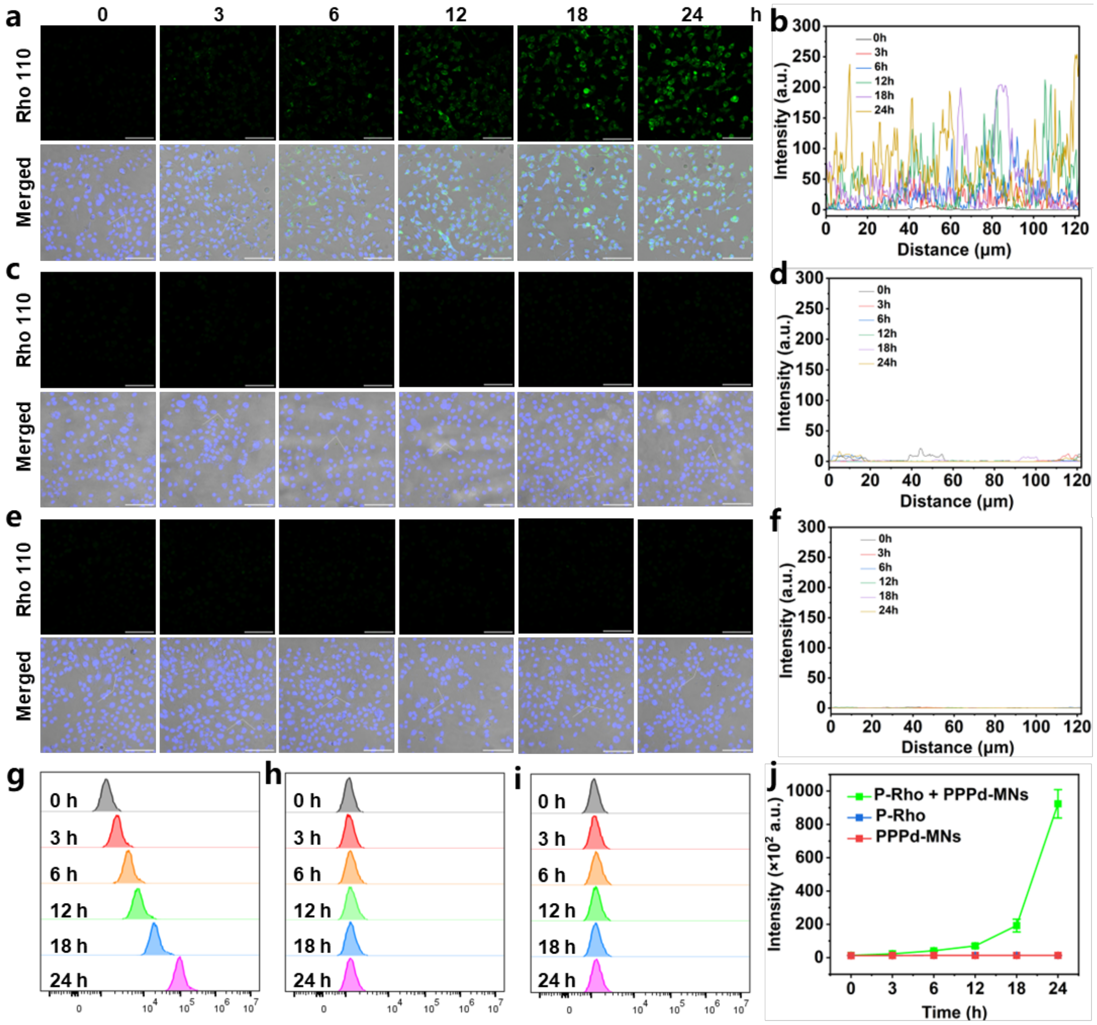


**Figure S10.** Intracellular bioorthogonal activation of P-Rho mediated by PPPd-MNs in 4T1 cells. (a–f) Representative confocal laser scanning microscopy (CLSM) images visualizing the time-dependent fluorescence recovery in 4T1 cells treated with the combination of P-Rho and PPPd-MNs (a, b), P-Rho alone (c, d), or PPPd-MNs alone (e, f). The corresponding line scan profiles (right panels) depict the fluorescence intensity distribution along the white dashed lines. Scale bar: 50 μm. (g–i) Flow cytometric histograms of 4T1 cells subjected to different treatments over time:

P-Rho + PPPd-MNs (g), P-Rho (h), and PPPd-MNs (i). (j) Quantified mean fluorescence intensity (MFI) derived from the flow cytometry data in panels (g–i). Data represent mean ± SD (n = 3).

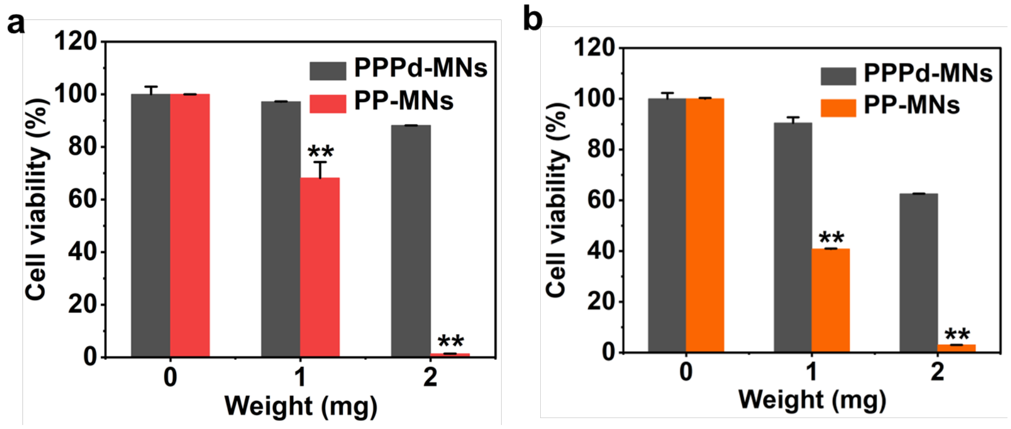


**Figure S11.** The cell viability of B16-F10 cells (a) and 4T1 cells (b) treated with PPPd-MNs or PP-MNs of different qualities for 48 h. Data points represent the mean ± standard deviation of three independent experiments (n = 3).

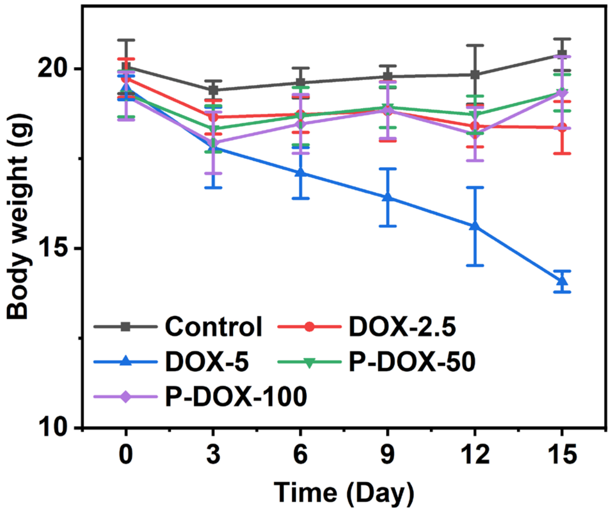


**Figure S12.** Assessment of systemic toxicity for DOX and P-DOX following intraperitoneal administration. Body weight fluctuations of mice were monitored throughout the treatment regimen (injections every 3 days). Notably, the group treated with DOX exceeding 5 mg $kg^{-1}$

exhibited severe weight loss, whereas P-DOX demonstrated a favorable safety profile with negligible toxicity. Data are shown as mean ± SD (n = 5 biologically independent mice).

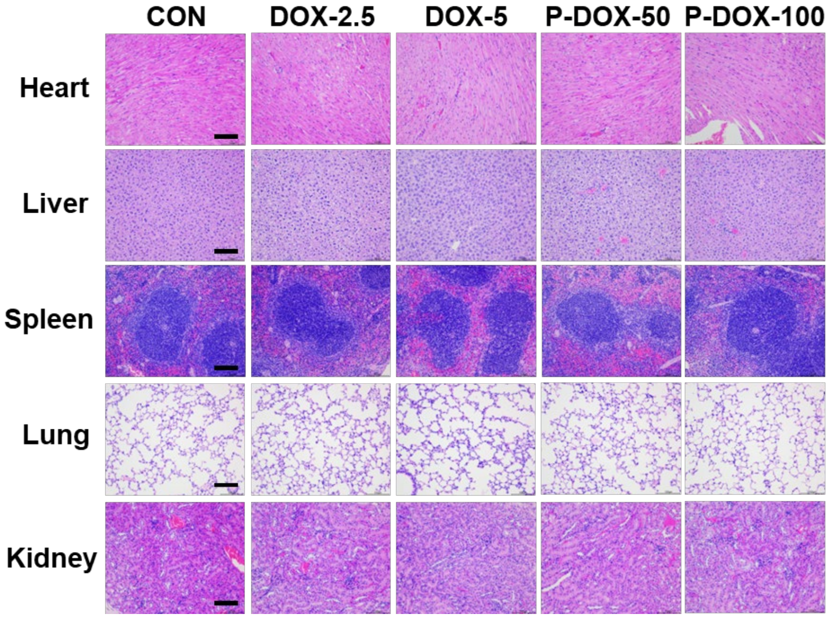


**Figure S13.** Histological examination of major organs harvested from mice after concluding different treatments (DOX, P-DOX, and PBS). Representative images of Hematoxylin and Eosin (H&E) stained sections are shown. Scale bar: 100 μm. Specific cardiac damage, characterized by myocardial structural disarray, widened interstitial gaps, and inflammatory cell infiltration, was observed in the high-dose DOX group (5 mg $kg^{-1}$).

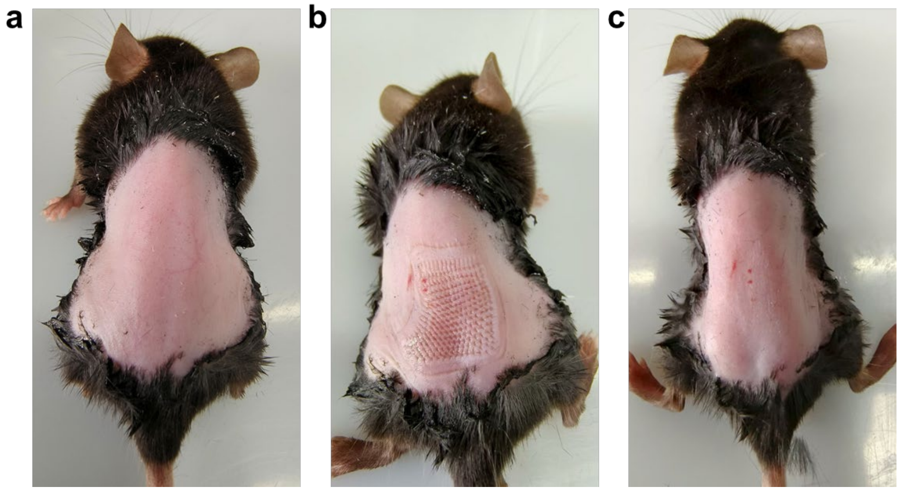

**Figure S14.** Skin insertion capability of the PPPd-MNs patches. (a) Photograph of mouse skin before a PPPd-MN patch insertion. (b) Micropores were observed after removal of the microneedle patch at the end of treatment. (c) The micropores were temporary and could be fully restored within 2 h after removing the PPPd-MNs.

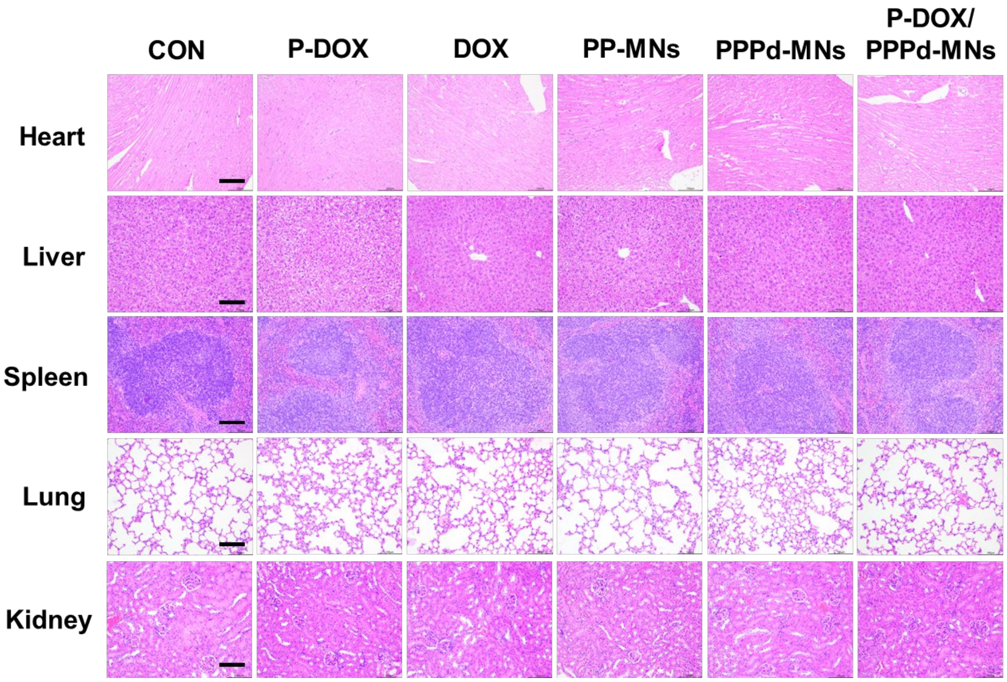


**Figure S15.** H&E staining images of representative major organs of mice after 12 days of different treatments, including PPPd-MNs, PP-MNs, DOX, P-DOX, P-DOX/PPPd-MNs and PBS. Scale bar: 100 μm.
